# PC-index, a composite metric for gene expression in single-cell RNA-seq

**DOI:** 10.64898/2026.01.21.700965

**Authors:** Ren Zhang

## Abstract

Single-cell RNA sequencing enables analysis of gene expression heterogeneity, but summarizing gene expression across large cell populations remains challenging. Gene expression is commonly described using average expression (counts per 10,000 transcripts, CP10K) and cellular prevalence (the fraction of cells expressing a gene), which can be difficult to interpret jointly. Here, we introduce the PC-index, a single, intuitive metric that integrates expression magnitude and prevalence. The PC-index is defined as the largest value X such that at least X% of cells express a gene at ≥X/10 CP10K. Using human adipocytes from GTEx single-nucleus RNA-seq as an illustrative example, we show that adipocyte genes exhibit distinct PC-index profiles. For example, adiponectin has a PC-index of 21, indicating that at least 21% of adipocytes express it at ≥2.1 CP10K. Conceptually, the PC-index reflects both percentage and CP10K, yielding a stable and interpretable single-number summary of gene expression behavior in single-cell data.

## Introduction

Gene expression profiles are routinely used to define cellular identity, infer regulatory and metabolic programs, and compare phenotypes across tissues, conditions, and individuals [1]. For decades, most transcriptomic analyses have relied on bulk measurements, which provide stable estimates of average expression but inherently mask cellular heterogeneity within complex tissues.

Single-cell and single-nucleus RNA sequencing extend gene expression analysis to cellular resolution, enabling systematic characterization of cell types, cell states, and gene expression variability within a population [2-4]. As these datasets grow in size and become increasingly integrated into atlases and disease studies, there is a practical need for summary measures that are both robust and interpretable, particularly when evaluating gene expression patterns across tens of thousands of genes within a defined cell population.

In routine single-cell analyses, gene expression within a cell population is often summarized using two complementary quantities. First, average expression (commonly reported as counts per 10,000 total transcripts, CP10K) captures expression magnitude. Second, cellular prevalence (the percentage of cells in which a gene is detected) captures how broadly a gene is expressed [4-6]. While informative, each quantity has limitations when considered alone. Average expression can be disproportionately influenced by a minority of high-expressing cells, obscuring whether expression is broadly distributed. Conversely, prevalence does not distinguish weak from strong expression and can assign similar weights to genes that are widely detected at low levels versus those expressed robustly in fewer cells. In addition, interpreting and comparing gene expression using two separate numbers complicates genome-wide ranking, marker selection, and cross-dataset comparisons.

Here we introduce the PC-index, a single, intuitive summary statistic that jointly captures expression magnitude and cellular prevalence in one interpretable value. Defined from the ranked distribution of per-cell expression, the PC-index increases only when expression is both sufficiently strong and sufficiently widespread across the cell population. Using human adipocytes from GTEx single-nucleus RNA-seq as an illustrative example, we show that PC-index provides a compact and easily interpretable representation of gene expression behavior, complementing commonly used measures of average expression and cellular prevalence.

## Materials and Methods

### GTEx single-nucleus RNA-seq data

Single-nucleus RNA-seq data were obtained from the GTEx v10 release [6]. Cell-type annotations provided by the GTEx consortium were used to define cell populations. Unless otherwise stated, analyses were performed using all cells annotated as adipocytes. Quality control and preprocessing followed GTEx consortium standards, and no additional cell-level filtering was applied. Gene expression levels were normalized as counts per 10,000 total transcripts (CP10K). A gene was considered expressed in a given cell if its CP10K value was greater than zero. For each gene, mean CP10K was calculated across expressing cells only, and cellular prevalence was defined as the percentage of cells in which the gene was detected.

### PC-index definition

Let C10 = CP10K × 10, a simple rescaling that places expression magnitude and percentage on comparable numeric scales. For each gene, cells are sorted in decreasing order of C10. The PC-index is defined as the largest value X such that at least X% of cells have C10 ≥ X.

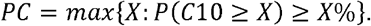

Here, P(C10 ≥ X) denotes the fraction of cells whose expression is at least X. Thus, PC = X indicates that at least X% of cells express the gene at an expression level of X or higher.

### PC-index calculation

For PC-index calculation, CP10K values were rescaled as C10 = CP10K × 10 and cells were ranked in decreasing order of C10 for each gene. Let N denote the total number of cells in the analyzed population. For a given expression threshold X, the fraction of cells with C10≥X was computed as k/N, where k is the number of cells meeting this criterion. Operationally, this was determined by iterating over ranked cells and identifying the maximum X for which at least X% of cells exhibited expression levels C10≥X. Genes with zero expression across all cells in the analyzed population were assigned a PC-index of zero.

### Statistical analysis, visualization and software

Associations between PC-index and mean CP10K expression were assessed using Spearman rank correlation. No additional hypothesis testing was performed unless otherwise specified. All analyses were conducted in Python (version 3.x) using Scanpy, NumPy, Pandas, and SciPy. Figures were generated using Matplotlib. Custom scripts were used for PC-index computation and data visualization.

## Results

### Gene expression heterogeneity in human adipocytes

As an illustrative example, we examined gene expression heterogeneity in human adipocytes using GTEx single-nucleus RNA-seq data. To characterize the joint behavior of expression magnitude and cellular prevalence, we plotted for each gene the percentage of adipocytes in which it is detected against its mean normalized expression level (CP10K) among expressing cells. This analysis revealed a highly constrained prevalence–intensity landscape (Figure 1A). The majority of genes clustered in a dense region characterized by low expression intensity and low cellular prevalence, whereas only a small subset of genes occupied a sparse upper frontier combining high expression with broad prevalence, corresponding to core adipocyte identity and functional programs.

**Figure 1.**
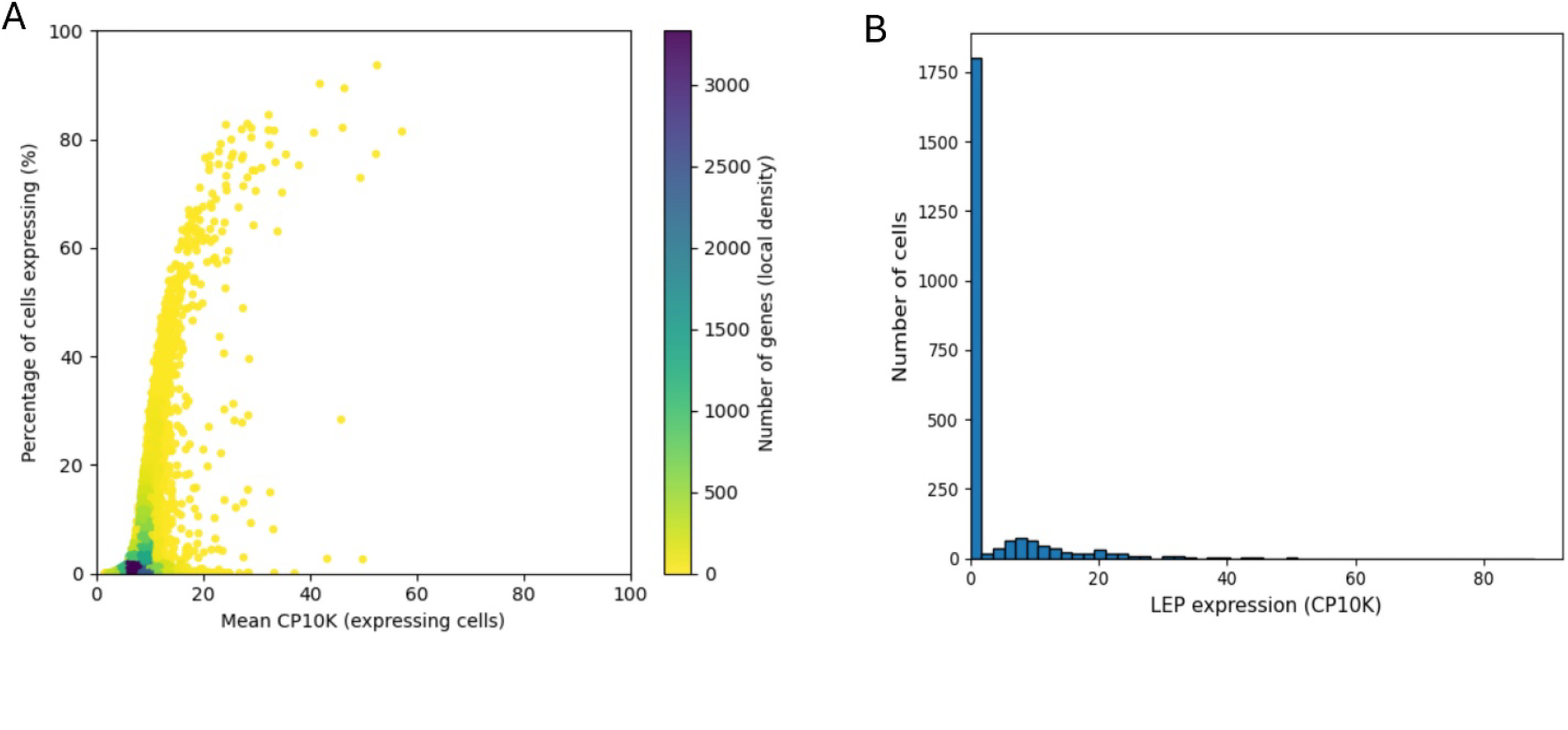
Gene expression landscape in human adipocytes. (A) Gene expression prevalence–intensity landscape. Each point represents a gene detected in GTEx single-nucleus RNA-seq adipocytes. Mean expression (CP10K) among expressing cells is plotted against the percentage of adipocytes expressing the gene; color indicates local gene density. The landscape is highly constrained, with most genes exhibiting low expression and low prevalence and a sparse frontier of genes combining high expression and broad prevalence, defining core adipocyte identity. (B) Distribution of LEP expression. Histogram of per-cell LEP expression (CP10K) in GTEx adipocytes. LEP expression is highly right-skewed, with most adipocytes lacking detectable transcripts and a minority exhibiting high expression, indicating pronounced cell-to-cell heterogeneity.

Leptin (LEP) is an adipocyte-secreted hormone that plays a central role in the regulation of body weight and energy homeostasis [7]. To further illustrate cell-level expression heterogeneity within this landscape, we examined the distribution of LEP expression across individual adipocytes. LEP expression exhibited a strongly right-skewed distribution, with most adipocytes lacking detectable transcripts and a minority displaying high expression levels (Figure 1B). This pronounced heterogeneity highlights that genes with similar average expression or detection frequency can differ substantially in how expression is distributed across cells.

Together, these observations suggest that gene expression in adipocytes is shaped by coupled but non-redundant dimensions of expression magnitude and cellular prevalence, motivating the development of a summary metric that integrates both properties, which we formalize as the PC-index.

### Definition of the PC-index

To integrate expression magnitude and cellular prevalence into a single summary metric, we defined the PC-index based on the ranked distribution of gene expression across cells (Figure 2). For each gene, cells are ordered by decreasing normalized expression (CP10K), and expression values are scaled as C10 (CP10K × 10). Plotting C10 against the cumulative percentage of cells yields a monotonically decreasing curve that captures how expression strength is distributed across the cell population. The PC-index is defined as the largest value X such that at least X% of cells exhibit expression levels C10 ≥ X, corresponding to the intersection of the ranked expression curve with the diagonal where expression equals cumulative cell percentage.

**Figure 2.**
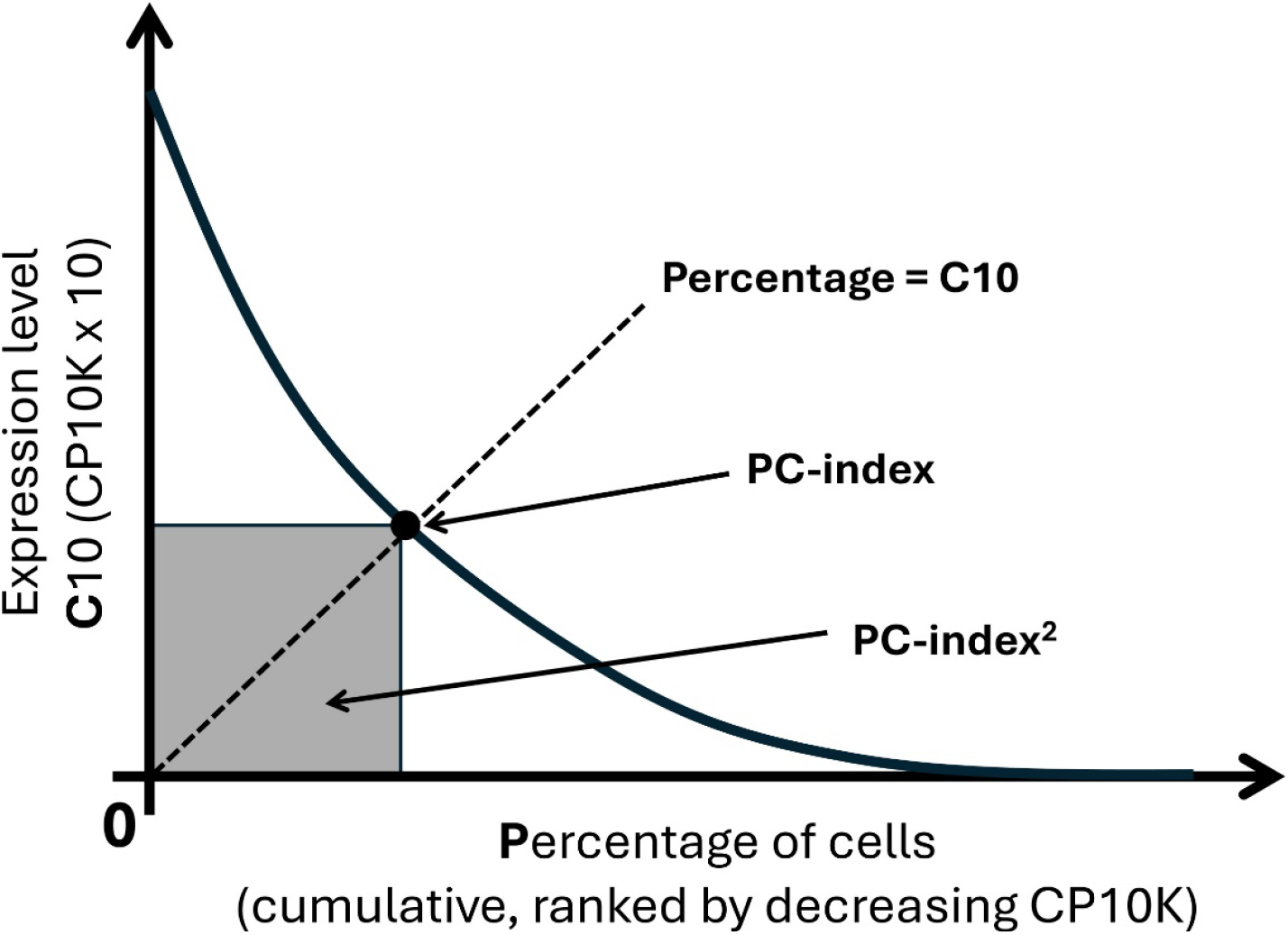
Definition of the PC-index. Cells are ranked by decreasing expression level measured as CP10K, and expression values are scaled as C10 (CP10K × 10). The x-axis shows the cumulative percentage of cells ranked by decreasing CP10K, and the y-axis shows C10 expression. The PC-index is defined as the largest value X such that at least X% of cells have C10 ≥ X, corresponding to the intersection of the ranked expression curve with the diagonal (C10 = cumulative percentage). The shaded square denotes (PC-index)^2^, representing the joint contribution of expression magnitude and cellular prevalence.

This construction identifies a fixed point that jointly reflects expression intensity and prevalence, assigning higher PC-index values to genes that are both strongly expressed and broadly detected, while down-weighting genes whose average expression is driven by either rare high-expressing cells or widespread low-level transcription. The shaded area, proportional to (PC-index)^2^, provides an intuitive geometric interpretation of the combined contribution of expression magnitude and cellular prevalence captured by the PC-index.

This behavior is illustrated by canonical adipocyte markers. PLIN1, a lipid-droplet protein essential for adipocyte lipid storage, exhibits a PC-index of 23.9, indicating that at least ∼24% of adipocytes express PLIN1 at a normalized expression level of ≥2.39 CP10K. Similarly, ADIPOQ, a broadly expressed adipokine characteristic of mature adipocytes, shows a PC-index of 20.9, corresponding to expression levels of ≥2.09 CP10K in approximately one-fifth of adipocytes. In contrast, LEP, an adipocyte-derived hormone, attains a PC-index of 10.9, indicating detectable expression of ≥1.09 CP10K in a smaller fraction of cells, despite a comparable mean expression level. Thus, PC-index quantitatively captures gene expression by jointly integrating expression magnitude and cellular prevalence, revealing differences in the breadth and consistency of expression across adipocytes that are not apparent from average expression alone.

### Genome-wide properties of the PC-index in adipocytes

We next examined the genome-wide distribution and expression dependence of the PC-index using human adipocytes as an illustrative example (Figure 3). Across all genes detected in GTEx adipocytes, PC-index values exhibited a strongly right-skewed distribution, with the majority of genes showing low PC-index values and a small subset attaining markedly high values (Figure 3A). This pattern indicates that only a limited number of genes combine substantial expression magnitude with broad cellular prevalence, consistent with the presence of a restricted set of core adipocyte identity genes.

**Figure 3.**
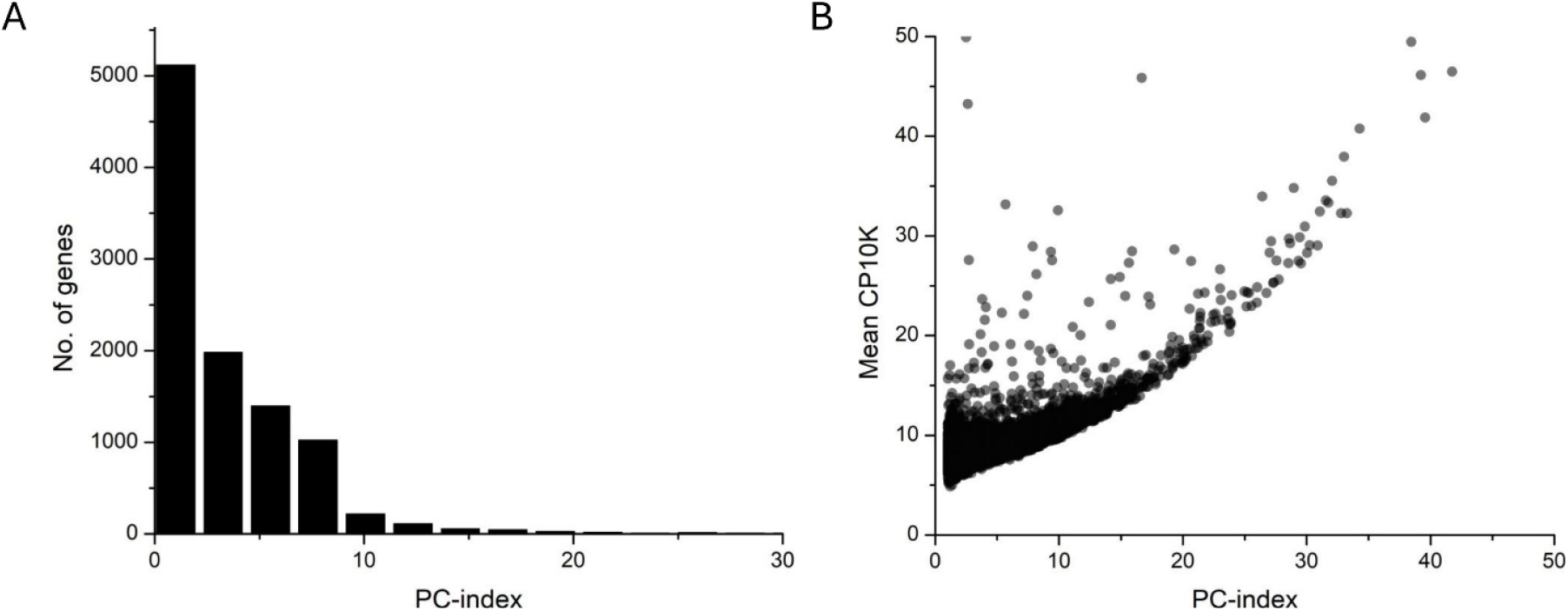
Genome-wide distribution and expression dependence of PC-index in adipocytes. (A) Distribution of PC-index values across genes expressed in GTEx adipocytes, showing a strongly right-skewed distribution in which only a small subset of genes attains high PC-index values. (B) Relationship between PC-index and mean gene expression (CP10K) in adipocytes. Each point represents a gene. PC-index shows a moderate positive correlation with CP10K (Spearman ρ = 0.63), indicating that PC-index is related to, but not determined by, average expression.

To assess the relationship between PC-index and average expression, we compared PC-index values with mean CP10K across genes (Figure 3B). PC-index showed a moderate positive correlation with mean expression (Spearman ρ = 0.63), indicating that genes with higher average expression tend to—but do not necessarily—exhibit higher PC-index values. Notably, substantial dispersion was observed at comparable expression levels, demonstrating that genes with similar mean CP10K can differ widely in PC-index depending on the fraction of cells in which they are expressed. Together, these results indicate that while related to expression magnitude, PC-index captures additional information by jointly integrating expression level and cellular prevalence, providing a complementary measure of gene expression behavior across adipocytes.

### PC-index prioritizes core adipocyte identity and metabolic programs

Ranking genes by PC-index in human adipocytes revealed a coherent set of genes with well-established roles in adipocyte identity, lipid metabolism, and hormonal regulation (Table 1). The top-ranked genes exhibited uniformly high PC-index values, reflecting the joint presence of substantial expression levels and broad cellular prevalence across adipocytes. Notably, these genes also showed high percentages of expressing cells, indicating that elevated PC-index values are characteristic of genes that are both robustly and consistently expressed within the adipocyte population.

**Table 1.** Top 20 adipocyte genes ranked by PC-index.

| Gene | PC-index | Mean CP10K | % Cells Expressing | Functional Category | Adipocyte-Specific Functional Summary |
| --- | --- | --- | --- | --- | --- |
| ACACB | 96.26 | 23.43 | 97.91 | Fatty acid metabolism | Controls malonyl-CoA levels to regulate fatty-acid oxidation and lipid storage in adipocytes |
| PTPRG | 91.87 | 18.31 | 93.49 | Adipocyte signaling | Modulates receptor-mediated signaling pathways involved in adipocyte metabolic regulation |
| EBF1 | 87.57 | 16.34 | 89.83 | Adipocyte lineage commitment | Transcription factor essential for adipocyte lineage specification and maintenance |
| FOXO1 | 86.00 | 16.64 | 88.98 | Insulin–adipocyte signaling | Integrates insulin signaling with lipid metabolism and adipocyte energy homeostasis |
| ACSL1 | 81.28 | 14.36 | 84.04 | Lipid activation & storage | Activates long-chain fatty acids for triglyceride synthesis in adipocytes |
| RBMS3 | 79.62 | 13.43 | 82.04 | Adipose stromal / structural program | Associated with mesenchymal and adipose stromal programs supporting adipocyte structure |
| COL4A2 | 79.28 | 13.54 | 82.43 | Adipose extracellular matrix | Basement membrane collagen shaping the adipocyte niche |
| COL4A1 | 78.51 | 13.26 | 81.53 | Adipose extracellular matrix | Structural collagen contributing to adipose tissue integrity |
| DMD | 78.30 | 13.67 | 81.28 | Adipocyte cytoskeleton | Supports membrane and cytoskeletal stability in lipid-laden adipocytes |
| PPARG | 78.13 | 13.69 | 81.11 | Master adipocyte regulator | Central transcriptional regulator of adipogenesis and lipid storage |
| LPIN1 | 78.00 | 15.28 | 81.57 | Triglyceride synthesis | Controls phosphatidate metabolism and triglyceride synthesis in adipocytes |
| FASN | 77.40 | 16.19 | 80.81 | De novo lipogenesis | Catalyzes fatty-acid synthesis supporting lipid accumulation in adipocytes |
| LINGO1 | 76.72 | 14.83 | 81.32 | Adipocyte membrane regulation | Membrane-associated protein reported in adipose-enriched regulatory contexts |
| FRMD4A | 76.47 | 12.6 | 79.57 | Adipocyte polarity / structure | Cytoskeletal adaptor contributing to adipocyte membrane organization |
| MLXIPL (ChREBP) | 76.38 | 13.22 | 81.91 | Glucose–lipid coupling | Glucose-responsive transcription factor driving adipocyte lipogenesis |
| BCL2 | 75.66 | 13.12 | 79.70 | Adipocyte survival | Promotes adipocyte survival during lipid loading and metabolic stress |
| GHR | 75.40 | 13.02 | 78.43 | Hormonal control of adiposity | Mediates growth hormone effects on adipocyte metabolism and lipolysis |
| SORBS1 | 74.72 | 12.02 | 76.89 | Insulin-responsive signaling | Adaptor protein supporting insulin-stimulated glucose uptake in adipocytes |
| MGST1 | 74.60 | 11.72 | 77.49 | Adipocyte redox homeostasis | Protects adipocytes from oxidative stress associated with lipid metabolism |
| SLC7A6 | 74.34 | 12.26 | 78.77 | Metabolic substrate transport | Amino-acid transporter supporting metabolic demands of adipocytes |

The highest-ranked genes were strongly enriched for canonical adipocyte metabolic pathways, including fatty acid synthesis and oxidation (e.g., ACACB, ACSL1, FASN, LPIN1), triglyceride storage, and glucose-lipid coupling (MLXIPL/ChREBP) [8,9]. Central transcriptional regulators of adipocyte differentiation and maintenance, such as PPARG, EBF1, and FOXO1, were also prominently represented, underscoring the ability of PC-index to recover core regulatory nodes defining adipocyte identity. In addition, genes involved in insulin and hormonal signaling (SORBS1, GHR), adipocyte survival (BCL2), and redox homeostasis (MGST1) were highly ranked, highlighting coordinated metabolic and stress-response programs intrinsic to mature adipocytes.

Beyond classical metabolic genes, PC-index also prioritized structural and microenvironmental components of adipose tissue, including extracellular matrix genes (COL4A1, COL4A2) and cytoskeletal or membrane-associated proteins (DMD, FRMD4A, LINGO1), reflecting the integrated structural and metabolic specialization of adipocytes. Collectively, these results indicate that PC-index effectively enriches for genes that define adipocyte identity and function by simultaneously capturing expression magnitude and cellular prevalence, rather than relying on average expression alone.

## Discussion

Single-cell RNA-seq data are inherently sparse and heterogeneous, making it challenging to summarize gene expression behavior using existing metrics alone. Average expression measures capture expression magnitude but can be disproportionately influenced by rare, high-expressing cells, whereas cellular prevalence reflects detection frequency without accounting for expression strength. Although interpreting these two quantities jointly is informative, it is often cumbersome in genome-wide analyses that require ranking or prioritization of thousands of genes. To address this challenge, we introduce the PC-index as a simple and intuitive summary metric for characterizing gene expression patterns in single-cell transcriptomic data. The PC-index integrates expression magnitude and cellular prevalence into a single, interpretable value, providing a compact description of how strongly and how consistently a gene is expressed across a population of cells. Unlike threshold-based marker definitions or heuristic score combinations, the PC-index is parameter-free and derived directly from the empirical expression distribution. It is designed to be easy to compute, interpret, and compare across genes and datasets, addressing a practical need in large-scale single-cell analyses.

The PC-index is intended to complement standard expression summaries such as average expression and cellular prevalence, rather than replace them. Its primary utility lies in situations where a single, intuitive metric is advantageous, including gene ranking, marker prioritization, cross-tissue comparisons, and exploratory analyses of large single-cell atlases. In these contexts, the PC-index provides a concise representation of gene expression behavior that is straightforward to communicate and interpret. Beyond its computational definition, the PC-index also has a natural conceptual interpretation: the abbreviation “PC” may be viewed as reflecting both percentage and CP10K, or equivalently prevalence and counts (per 10,000 transcripts). This dual interpretation emphasizes that the metric jointly captures how many cells express a gene and at what level, reinforcing its intuitive appeal and facilitating its adoption without requiring detailed familiarity with its mathematical formulation.

Using human adipocytes as an illustrative example, we show that the PC-index naturally prioritizes genes with well-established roles in adipocyte identity and metabolism. Canonical adipocyte markers with robust and widespread expression attain high PC-index values, whereas genes with heterogeneous or sparse expression exhibit lower values despite comparable average expression levels. These patterns are consistent with known adipocyte biology, underscoring the interpretability of the PC-index and its ability to capture biologically meaningful expression behavior without reliance on cell-type-specific tuning or arbitrary thresholds.

Several limitations should be acknowledged. PC-index values depend on data quality, normalization strategy, and sequencing depth, and extremely sparse datasets may yield uniformly low values. In addition, by collapsing expression magnitude and cellular prevalence into a single summary statistic, the PC-index necessarily entails a loss of information and does not capture the full distribution of expression values across cells. As a result, genes with distinct underlying expression patterns may share similar PC-index values. Furthermore, the PC-index summarizes expression within a defined cell population and does not capture temporal dynamics, regulatory interactions, or cell-cell communication. These limitations should be considered when interpreting PC-index values in specific biological contexts.

Looking forward, the PC-index may be useful for prioritizing genes in single-cell studies, including the identification of cell-type markers, comparison of gene expression programs across tissues or conditions, and integration with downstream analytical frameworks. Although we focus here on adipocytes as an illustrative example, the PC-index is defined independently of cell type, tissue, or species and can be applied to any single-cell or single-nucleus transcriptomic dataset in which per-cell expression distributions are available. Future work may explore extensions of the PC-index to alternative normalization schemes, its behavior across diverse cell types and datasets, and its incorporation into differential expression or atlas-scale analyses.

In summary, the PC-index provides a practical, intuitive, and interpretable approach for summarizing gene expression in single-cell transcriptomic data. By jointly integrating expression magnitude and cellular prevalence into a single number, it offers a useful complement to existing measures and facilitates clearer and more efficient analysis of gene expression heterogeneity.

## Notes

### Competing Interest Statement

The authors have declared no competing interest.

